# HUMESS: Integrating Quantitative Transcriptomic Analysis and Metabolic Modeling to Unveil Condition-Specific Gene Signatures

**DOI:** 10.1101/2025.02.28.640738

**Authors:** Louis Paré, Philippe Bordron, Laurent David, Maxime Mahé, Audrey Bihouée, Damien Eveillard

## Abstract

Transcriptomic analysis is a key tool for exploring gene expression, but the complexity of biological systems often limits its insights. In particular, the lack of intermodal or multi-layered analysis hinders the ability to fully capture key cellular functions such as metabolism from transcriptomic data alone. Here, we introduce a novel approach that integrates transcriptomic data with metabolic network modeling to address this. Unlike traditional methods, HUMESS prioritizes genes based on their metabolic significance, offering a deeper understanding of condition-specific gene expression. Our computational pipeline, supported by a user-friendly Rshiny application, enhances gene expression analysis by uncovering metabolic phenotypic signatures.

## Introduction

Transcriptomic analysis is a powerful tool for elucidating gene expression patterns associated with specific biological conditions, offering invaluable insights into cellular responses and regulatory mechanisms (1). However, despite its utility, many transcriptomic studies fail to provide comprehensive insights due to the inherent complexity (i.e., multilayered omics intricacy) of biological systems and the limitations of purely expression-based approaches. One major challenge in transcriptomic analysis is the need for external knowledge to interpret gene expression changes in a meaningful biological context, which is labor-intensive and prone to biases (2). Consequently, many gene expression signatures derived from transcriptomic data remain superficial, and lack the depth necessary for proper mechanistic understanding. Whereas conventional methodologies perceive alterations in gene expression in isolation from one another, contemporary approaches endeavor to consolidate genes into extensive collections for the evaluation of phenotypic characteristics. For example, network analyses advocate for the aggregation of genes predicated on their co-expression (3) or the association of genes based on their mechanistic activation or repression (4). More recently, initiatives have been directed towards quantitative biology to evaluate metabolic networks. These networks exemplify a collection of reactions that are encoded by gene products and are interrelated when the output of one reaction serves as the substrate for another. Recent advancements have facilitated the conversion of these networks into genome-scale metabolic models (i.e., GEMs) to evaluate the fluxes executed by all metabolic reactions(5). The assessment of such quantitative insights encapsulates metabolic phenotypes and is particularly applicable to human data, as it encompasses the most comprehensive dataset of multi-modal information. Nonetheless, despite the emergence of context-specific metabolic models that are classified as Human GEMs(6), the integration of diverse biological data continues to pose a significant challenge, as it necessitates the harmonization of information from multiple sources to accurately reflect the intricacies of human metabolism. Furthermore, these GEMs are infrequently used to inform analyses of gene expression.

Here, we introduce a novel approach termed HUMESS (HUman MEtabolism Specific Signature), which seeks to bridge this gap by leveraging recent advances in quantitative transcriptomic analysis and metabolic modeling. Unlike traditional methods that treat gene expression changes in isolation, HUMESS integrates quantitative transcriptomic data with metabolic network modeling to unveil condition-specific gene signatures with enhanced biological relevance. The originality of this work lies in the use of methods initially developed for modeling bacterial ecosystems on human metabolism (7). By combining transcriptomic analysis with metabolic modeling, HUMESS offers a unique framework for identifying genes that are not only differentially expressed under specific conditions but also play critical roles in shaping specific human metabolic phenotypes. This integration allows for the prioritization of genes based on their metabolic significance, providing a more nuanced understanding of cellular responses and functional implications.

This manuscript describes a computational pipeline that uniquely combine RNA data and metabolic model reconstruction in Python to perform analysis. For a user friendly interface, the pipeline is accompanied by a Rshiny web application (8) that permits loading pipeline output files for manipulating various graphical representations (i.e., MA and Volcano plots). We demonstrate the utility of HUMESS by improving the analysis of recent case studies in human development (9). By elucidating condition-specific gene signatures weighted by their metabolic relevance, we showcase the power of integrating quantitative transcriptomic analysis and metabolic modeling to uncover novel insights into biological systems. Through HUMESS, we aim to provide researchers with a valuable tool for holistically and meaningfully exploring gene expression data, ultimately advancing our understanding of complex biological phenomena.

## Results

### Combining transcriptomic analysis and Human metabolic modelings

HUMESS has been developed using the 3’SRP pipeline for transcriptomic analysis as input (10). In contrast to conventional RNA-seq, 3’SRP analysis has the advantage of multiplexing samples and barcoding transcripts with single molecule identifiers (UMIs), significantly improving the accuracy of mRNA quantification. The HUMESS pipeline embeds gene-expressed data within a metabolic modeling framework for reconstructing a metabolic network of the human systems analyzed. Beyond the sole availability of the tuned metabolic model, HUMESS allows its systematic exploration for emphasizing features such as essential reactions that will enrich the transcriptomic analysis with phenotypic biomarkers. The pipeline follows the steps below illustrated in Fig. 1A.

**Fig. 1.**
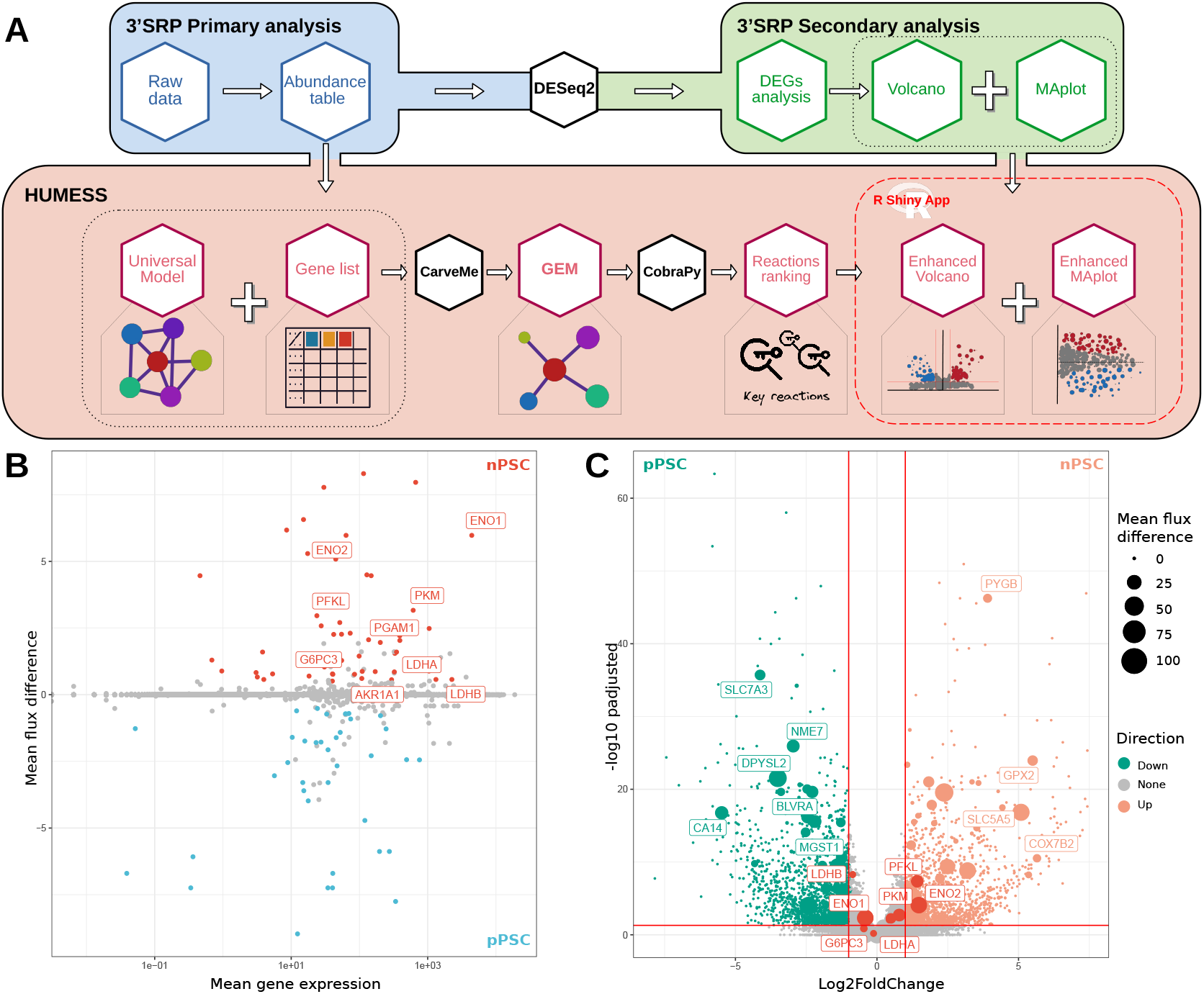
Description of the HUMESS pipeline and associated representations. A. HUMESS feeds on standard outputs of transcriptomic analysis to build a GEnome-scale Metabolic (GEM) model via a top-down approach. Sampling analyses are then performed to identify the reaction cumulative correlation score (RCC) and mean flux per GEM’s reactions. B. Projection of mean gene expression against the differential of mean flux values of associated metabolic reactions between the nPSC and pPSC conditions. C. Volcano plot comparing the nPSC and pPSC conditions with the size of nodes proportional to the absolute value of the difference of mean fluxes between conditions.

### Reconstruction of Human metabolic models from expressed genes

The rationale of this reconstruction consists of considering a universal metabolic model that embeds all available metabolic knowledge (i.e., a set of biochemical reactions satisfying the mass action law and thermodynamic constraints), and to build a context-specific metabolic model (i.e., for each experimental condition). HUMESS uses UMI count tables to feed an adaptation of a top-down metabolic network reconstruction, called CarveMe(7). This protocol consists of *carving* the universal metabolic model to fit genes expressed in the given condition while maintaining a functional overall metabolic model (i.e., keeping fluxes metabolic to produce essential metabolites for biological survival). Initially developed on prokaryotes, we improved CarveMe to (i) use a genome-scale human meta-model, called Recon3D(11), obtained from the BIGG(12), a database that contains high-quality, manually-curated GEMs; and (ii) use a list of expressed genes instead of a genome (see HUMESS documentation for details). Adaptation of CarveMe allowed the estimation of a Human metabolic model per condition that embeds all significantly expressed genes.

### Exploration of the metabolic model solution space

We can analyze reconstructed metabolic models via the COBRApy library (13). In HUMESS, we advocate for systematically explore the solution space of each human genome-scale model using randomized flux sampling analysis. For this purpose, HUMESS embeds two complementary techniques: a global solution space sampling via OptGPSampler(14) and the corner-based sampling (15). The first technique emphasizes the phenotype in exploring the whole solution space. This is computationally challenging, especially for large numbers of samples and a high thinning rate (i.e., reducing redundant data points). The second approach focuses on exploring the boundaries (corners) of the solution space. Therefore, this exploration weighs more quickly on extreme behaviors of the metabolic phenotype. The limits of sampling the solution space are undersampling (i.e., the lack of convergence) or oversampling (i.e., magnified autocorrelation). HUMESS avoids these limitations by using Raftery & Lewis and Geweke’s convergence tests (16) to find the optimal number of samples for exploring solutions. Sampling procedures generate a large tab-delimited file containing the flux values of all reactions of a given model, for each sampled points.

### Identification of essential reactions and associated genes

Sampling results are then processed via pairwise correlation between the reaction’s fluxes. On this basis, we compute the pairwise correlation of each metabolic reaction with every other reaction throughout all samples, resulting in a symmetric correlation matrix. We state each reaction’s importance (i.e., Reaction Cumulative Correlation - RCC) by summing up its absolute correlation values, then normalized to a scale of 0 to 100. Thus, a reaction associated with a RCC of 100 which is highly correlated with all given GEM reactions, is considered essential to sustain metabolic phenotypes. A distribution of RCC (i.e., histogram) or a list of all reactions ranked based on RCC assesses a comparison between different metabolic models. Furthermore, the mean value of all fluxes (*mol.kgDW* ^*−*1^.*h*^*−*1^) sampled per reaction is calculated. Worth noting, mean value of all metabolic fluxes reflect the reaction usage (i.e., amount of metabolic material passing through), whereas RCC reflect the importance of the reaction in maintaining the metabolic phenotype (i.e., weighted centrality).

Using the Gene-Protein-Reaction rules obtained from the BIGG database, RCC and usage of reactions is mapped to the genes involved in those reactions. HUMESS incorporates these scores in various state-of-the-art visualizations. An MA-plot-like visualization (see Fig. 1B) describes each GEM’s gene expression level compared to its associated reaction importance score. Similarly, conditions are compared via Volcano plots enriched with the differential reaction importance score between two given GEMs, represented as node size (see Fig. 1C).

Visualizing and ranking genes based on the importance of their associated reactions allows for the selection of genes for further analysis (Fig. 1B). This process can be applied to enrichment analyses, such as Over-Representation Analysis (ORA) using Gene Ontology (GO) terms or KEGG path-ways, or to Gene Set Enrichment Analysis (GSEA) (17), using the difference in reaction importance as a discriminatory score.

### Magnifying the whole stem cell transcriptomic analysis

To validate our approach, we applied HUMESS to existing transcriptomic data from various stages of human embryonic stem cell development. Specifically, Onfray and colleagues (9) examined several states of pluripotency : (i) naive (nPSC, modeling human epiblast between six to nine days after fertilization), (ii) primed pluripotent stem cells (pPSC, modeling human epiblast between ten to fourteen days after fertilization), (iii) extended PSCs (ePSC, pluripotent stem cells with extended potential) and (iiii) trophoblast stem cells (TSC, derived from human blastocyst but also from first trimester placenta). Authors discovered thousand of genes showing differential expression between all these states.

Building on these findings, we developed four types of GEMs for stem cells: the nPSC-like GEM (2,087 metabolites, 2,933 reactions and 509 genes), the pPSC-like GEM (1,961 metabolites, 2,692 reactions and 508 genes), the ePSC-like GEM (1,900 metabolites, 2,583 reactions and 484 genes), and the TSC-like GEM (2,495 metabolites, 3,773 reactions and 618 genes). Flux sampling was carried out with a thinning value of 10,000 for each of the four GEMs, resulting in sample sizes of 100,154 for nPSC, 104,670 for pPSC, 150,000 for ePSC, and 105,517 for TSC.

We used HUMESS to improve the interpretation of two comparisons. Firstly, we focused on the nPSCs compared to the pPSCs conditions, in which the metabolic differences have been widely studied (18). Specifically, we focused on significantly expressed genes (i.e., p-value < 0.05) exhibiting modifications in associated reaction flux (i.e., above 0.5 for nPSC vs. pPSC and below -0.5 for pPSC vs. nPSC). In the nPSC vs pPSC scenario, KEGG pathway enrichment analysis revealed that the genes most affected by alterations in metabolic flux were linked to glycolysis and glyconeogenesis pathway, highlighting 11 genes (ENO1, ENO2, PFKL, PGAM1, PKM, G6PC3, AKR1A1, LDHA, LDHB) even though only 2 of them exhibit a significant expression fold change (> one absolute Fold Change) (see Fig. 1C). Furthermore, the fourth most enriched pathway is the biosynthesis of amino acids. Together, these findings support the metabolic distinctions pointed out by Gu et al., highlighting an increase in glycolysis in the nPSC condition; their metabolomic study also highlighted an increased use of glucose for the biosynthesis of nucleotides and serine. The strength of HUMESS resides in elaborating a specific metabolic network for each condition, which can be used to deepen the analysis of the differences between the two conditions, notably by highlighting exactly which reactions are responsible for the shown difference. Here, we could discriminate the phosphoglucomutase reaction, responsible for converting glucose-1-phosphate to glucose-6-phosphate at the very start of the glycolysis process, with a higher metabolic flux in nPSC against pPSC. This reaction is linked to the PGM2 gene, preferentially expressed in nPSC. Two other reactions were also found with higher flux in nPSC towards the end of the glycolysis pathway, the enolase and the pyruvate kinase reaction, which are responsible for the generation of pyruvate. The ePSC vs pPSC comparison was also studied using HUMESS; the transcriptomic analysis by Onfray et al. showed only 180 differentially expressed genes. Using HUMESS, we could discriminate genes that exhibit higher reaction flux in the ePSC conditions and search for enriched pathways. The third most enriched pathway is oxidative phosphorylation, while glycolysis/gluconeogenesis is among the enriched pathways. Both of those findings support the metabolic distinctions pointed out by Onfray and colleagues, demonstrating the significance of the extracellular acidification rate directly associated with glycolysis and the oxygen consumption rate linked to oxidative phosphorylation to discriminate the two cell types.

## Conclusions

HUMESS enhances transcriptomic analysis as an independent tool by pinpointing significant genes that expression data alone may overlook. Beyond ranking the gene set, we propose that HUMESS could also inform metabolomics studies by identifying a collection of metabolites anticipated to be produced in excess within particular GEMs. Viewed from a wider lens, the primary aim of HUMESS is to recommend specific GEMs suited for future mechanistic research. We anticipate that this will aid in uncovering genes that maximize specific metabolites and allow for more thorough investigations into the transitions among various cell states, including those related to cell differentiation, as well as the exploration of the entropy associated with biological systems.

## Code availability, deployment and reproducibility

HUMESS is open source and available under GitLab (https://gitlab.univ-nantes.fr/bird_pipeline_registry/humess). Analyses were conducted on a cluster comprising 96 processors and 384 Go memory. We performed analysis using the CPLEX solver, which is free of charge on the IBM website for academic users, but HUMESS considers others open source solvers. An RShiny application has been developed to facilitate the exploration and analysis of HUMESS’s results. The app is available online at the following address: https://shiny-bird.univ-nantes.fr/app/Shinymess.

## Author contributions statement

L.P., A.B. and D.E. conceived the experiment(s), L.P. and P.B. conducted the experiment(s), L.P. and L.D. analyzed the results. L.P., A.B. and D.E. wrote and all authors reviewed the manuscript.

## Acknowledgments

This work was supported in part by funds from Pays de la Loire region via the GIS Biogenouest. The analyses and hosting of the shiny application are supported by the Bioinformatics Core Facility BiRD, member of Biogenouest and Institut Français de Bioinformatique (IFB) (ANR-11-INBS-0013) and by GLiCID (Groupement Ligérien pour le calcul Intensif Distribué, www.glicid.fr) computing resources.

